# Specific Fabric Properties Elicit Human Neuro and Electrophysiological Responses

**DOI:** 10.1101/2023.11.24.568599

**Authors:** Mahendran Balasubramanian, Thamizhisai Periyaswamy

## Abstract

Tactile perception of fabrics is a complex process challenging to parse and understand. Most studies examined fabric tactile perception through subjective means. Alternatively, neural and electrodermal signals can be used to examine fabric properties that elicit responses via tactile stimulation. Among the fabric properties, some may generate strong responses while others produce weaker responses. Here, we have shown that by studying the electroencephalography signal from the scalp areas corresponding to the sensory cortex, and the galvanic skin responses that occurred during active tactile sensation of varied fabrics, the influence of individual fabric properties can be determined. Eight subjective properties, from twenty female subjects, and seventeen objective properties, measured using Kawabata Evaluation System, were independently correlated to the biosignals. We have shown that properties related to thickness and volume were the most influential, followed by surface frictional properties. Fuzziness and linearity of compression were the least influencing properties affecting the biosignals.

## Introduction

When humans touch and feel a piece of fabric, they perceive the stimuli holistically and arrive at a characteristic notion of its properties. It is easy and quick for humans to perceive a fabric. Nevertheless, measuring the perception is challenging. And this is the primary reason fabric tactile comfort studies are primarily subjective. Complex processes occur in the human body while evaluating the tactile characteristics of materials, i.e., (i) physical properties of the materials, providing external stimuli to the skin, (ii) physiological and neurological responses in the body reacting to the external stimuli and (iii) cognitive psychological process formulating the overall subjective perception (McGlone and Spence, 2010). Of these three components, the physical properties of fabrics are understood through their mechanical behavior, i.e., tensile, bending, compression, shear, and surface properties. Measurement systems like Kawabata Evaluation System (KES) and Fabric Assurance by Simple Testing (FAST) measure these physical and mechanical properties at low-stress levels (Giorgio Minazio, 1995; Kawabata and Niwa, 1996). The third component, the cognitive psychological perception of fabric, has also been extensively studied with subjective assessment of tactile descriptors such as softness, stiffness, and roughness (Cardello *et al*., 2003; Mahar *et al*., 2013; Sülar and Okur, 2007). Its relationship with the physical properties of fabrics is also increasingly understood (Bertaux *et al*., 2007; Chen *et al*., 2009). The intermediate and perhaps the most informative component, i.e., the biosignals emanating from the human body during tactile stimuli, has received less attention, and its physiological and neurological mechanism is less understood.

Recently, studies have attempted to establish human neurological responses to fabric tactile stimuli. Neural functional areas and their responses to fabric touch were investigated using functional Magnetic Resonance Imaging (fMRI) (Wang *et al*., 2016, 2019; Yuan, Yu, Chen, *et al*., 2019) and electroencephalography (Hoefer *et al*., 2016; Jiao *et al*., 2020). They reported activations in the regions of the somatosensory cortex, i.e., S1 and S2 of the cortical areas. Another study by Rajaei et al. also reported the somatosensory cortex (S1) association with the tactile perception of velvet fabric textures. In addition, cortical sensory response to different fabric stimuli showed that neural activation information could distinguish the materials (Wang *et al*., 2019). Other studies have investigated brain activation related to pleasantness in fabrics and other textured materials (Faucheu *et al*., 2019; Singh *et al*., 2014). The focus of the existing studies was predominantly on identifying which brain areas were active during touch and how they varied with the type of fabric or texture. For example, Wang et al. studied brain regions responsible for tactile perception when a piece of fabric, either silk or linen, is touched between the thumb and index fingers of the right hand. These studies addressed the gross impact of fabric tactile stimuli on neural responses. The critical question of to what extent the fabric properties contribute to a given holistic perception during touch was left unexplored.

Furthermore, the existing studies use a passive touch, i.e., a fabric can rest on skin or be stroked over an open region or slid through digits in a pinch posture. However, it is critical for fabric hand evaluation to use active touch, i.e., feeling fabrics freely between the digits and the palm, as it affects how tactile information is processed and integrated. Besides, no reliable metrics or measurement techniques were established for bio-sensory analysis of textile materials in these studies. These obstacles limit the usage of physiological and neurological components in the objective evaluation of fabric hand.

In the present study, we explored the question of how well each fabric property, evaluated subjectively and objectively, correlated to two important bio signals to determine the most and least influential fabric properties on human perception. We measured the electroencephalography (EEG) and the galvanic skin response (GSR) signals during an active touch paradigm, and estimated their relationship with fabric properties. We used active touch, as it closely resembles a natural interaction paradigm. We showed that specific fabric properties are highly correlated to the α-band (8-14 Hz) in EEG, corresponding to attention in the somatosensory cortex and the number of fluctuations in the GSR.

## Material and Methods

### Subjects

Twenty volunteer female subjects with a mean age of 20.5 years (± S.D. 1.5 years) participated in the study. Since this study involved physiological recordings, subjects undergoing medical treatments or medications/drugs were excluded. All the subjects were right-handed and received training on making more uniform contact with fabrics using their fingers and palm without moving the forearm. Participants signed the informed consent approved by the Institutional Review Board of the host institute.

### Stimuli

A pool of twenty-five fabrics was tested using the Kawabata Evaluation System (KES) to determine their LSM properties. KES is a standardized system for analyzing tactile characteristics of the fabric and has been extensively used to understand the hand values of various fabrics (Kawabata and Niwa, 1996). Based on the KES results, six specimen fabrics were chosen for this study that represents a range of LSM properties. The fabrics were carefully chosen to represent a range of stiffness, roughness, and other physical attributes. A sample size of 10×10 cm was used in the experiment to touch and feel the fabric.

### Experimental Procedure

#### Setup

In a special hand evaluation booth (see Figure-1a), subjects were seated on one side, with sensors to measure EEG and GSR hooked to them, and the fabric swatches were presented from the other side of the booth. The opaque partition restricted the subjects from having any visual feedback of the fabric. The subjects were seated in a relaxed condition, with their right forearm resting on the booth table. They were allowed to sense the fabrics freely using their fingers over the palm area without moving their forearm. Each subject was asked to touch and feel the six different fabric samples. An assistant on the other side of the booth presented the fabrics to the subjects for evaluation. The samples were randomized prior to the presentation. The participants felt the fabrics with a normal contact force and were trained before the experiment on making regular contact with samples. Simultaneously, the sensors measured the GSR and EEG activity of the subjects.

**Figure 1.**
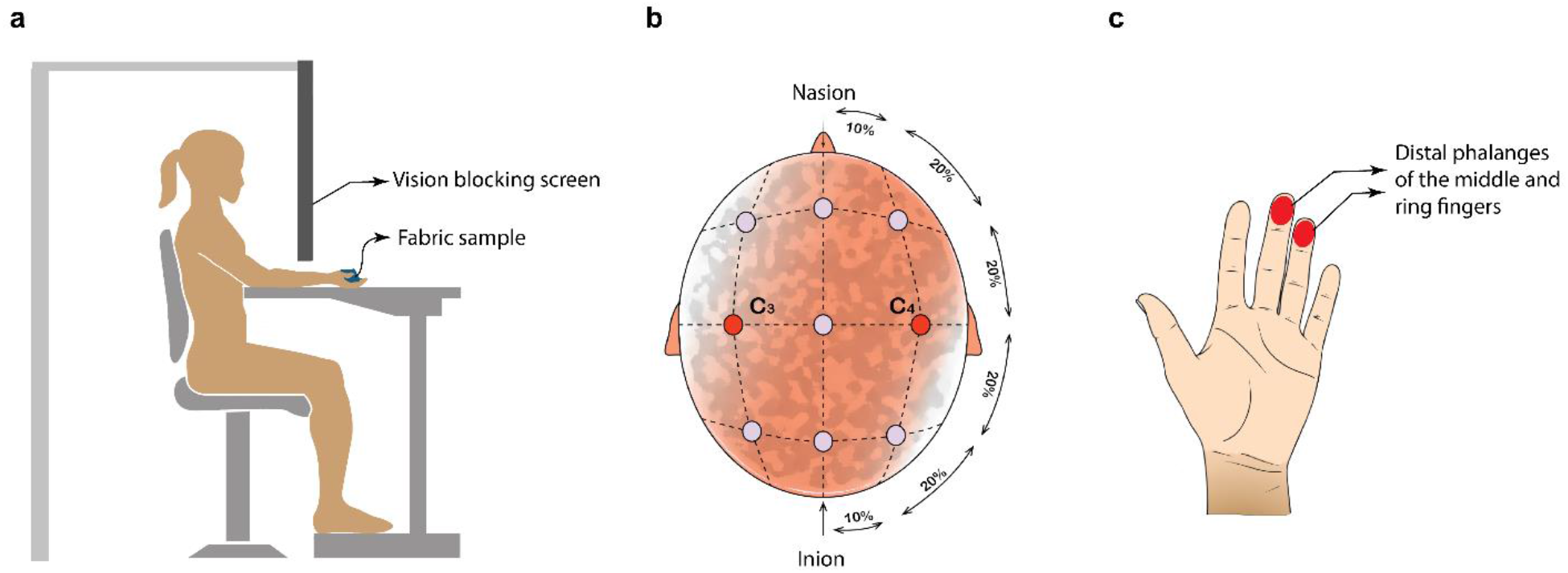
(a) booth set-up for fabric tactile sensation, (b) schematic of a montage showing the scalp locations C3-C4 chosen for EEG signal acquisition and (c) the distal phalanges of human finger chosen for EDA signal acquisition.

The recordings were segmented into three sessions, where the first and last sessions correspond to the pre and post-baseline recordings. The second session corresponded to the actual stimuli presentation. The pre and post-baselines were identical to the experimental conditions, except that the tactile stimuli were absent. Baselines were recorded for two minutes right before and after the tactile stimuli. The primary purpose of the baseline is to allow us to extract the physiological responses primarily caused by an externally applied tactile stimulus, excluding individual variance in the sample and other environmental conditions.

### Neuro and Electrophysiological Signals

Two electrodes mounted on the subjects’ scalps in the C3-C4 areas (see Figure 1b for a representative montage) that correspond to sensory regions of the brain were used to record the EEGs. In addition, galvanic skin responses emanating from the skin were measured using skin conductivity sensors mounted on the middle and ring fingertips of the left hand (see Figure 1c). We took extra caution while recording the EEGs to prevent (a) electrode and cable movements causing motion artifacts and (b) masticatory muscle tension causing electromyography artifacts. Outputs from the sensors were acquired in real-time via a clinical-grade neuro-biofeedback system (Nexus 10; Mindmedia Inc.) and stored for offline analyses. The EEG signals were sampled at a rate of 256 Hz and the GSR at 32 samples/second for 30 seconds per fabric sample. The signals were analyzed using Matlab 2022a.

### Subjective Evaluation

Following the experiment, the subjects immediately completed a subjective hand evaluation form for each fabric sample. The subjective fabric hand attributes (see Table-1) were adapted from Cardello et al. (Cardello *et al*., 2003) and modified to fit with a 0-9 Likert scale.

**Table 1.** Subjective Hand Evaluation Attributes.

| Properties | Definition |
| --- | --- |
| **Grainy**<br>[0-not grainy; 9-highly grainy] | Amount of small, round particles in the surface of the sample |
| **Gritty**<br>[0-not gritty; 9-highly gritty] | Amount of small, abrasive, picky particles in the surface of the sample |
| **Fuzziness**<br>[0-not fuzzy; 9-highly fuzzy] | Amount of pile, fiber, fuzz on the surface of the sample |
| **Thickness**<br>[0-very thin; 9-very thick] | Perceived distance between the thumb and index fingers (when the sample is placed between the two) |
| **Hand friction**<br>[0-nohand friction; 9-high hand friction] | Force required to navigate the digits across the surface of the sample |
| **Fabric-fabric friction**<br>[0-no fabric-fabric friction; 9-high fabric-fabric friction] | Force required to move the fabric over itself |
| **Stiffness**<br>[0-not stiff; 9-very stiff] | Degree to which the sample feels pointed, ridged, and cracked; not pliable |
| **Fullness/volume**<br>[0-not full; 9-very full] | Amount of material felt in the hand |

## Results

### Low-stress Mechanical Properties of the fabrics

The LSM properties of the fabrics used in this study are listed in Table-2. Fabric F1 and F2 were the lightest fabrics, 108 g/m^2^, and F6 was the heaviest, 278 g/m^2^. Out of the six fabrics, F6 showed the highest magnitude for 11 fabric properties, and F1 showed the lowest for six of the fabric properties. The remaining fabrics had magnitudes distributed in between. The mean LSM values of the fabrics are shown in Figure-2a.

**Table 2.**
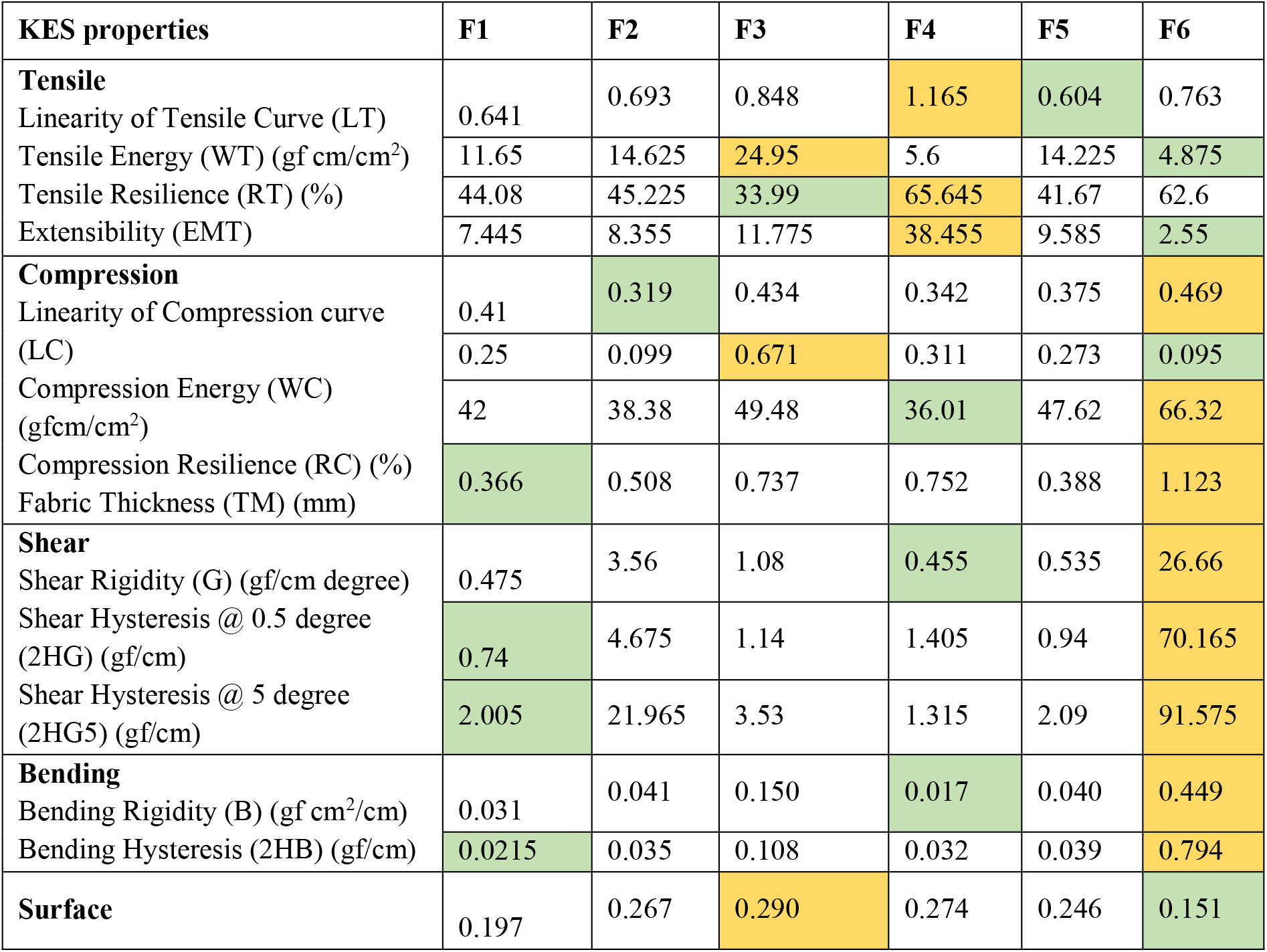
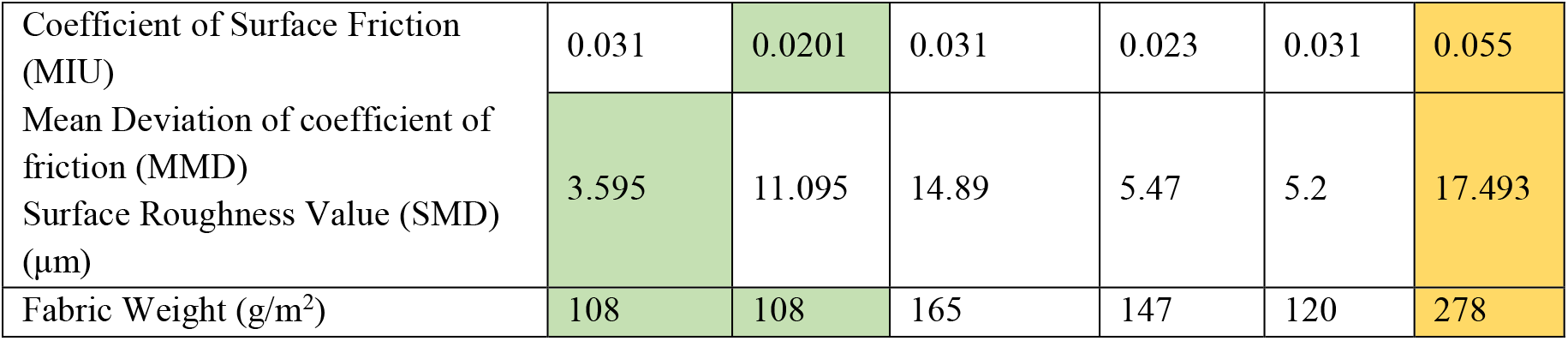
Low Stress Mechanical Properties of the six fabric samples (F1-F6) measured using the Kawabata Evaluation System (KES). Green filled boxes indicate the minimum magnitude for a given property and the orange colored boxes correspond to the maximum.

**Figure 2.**
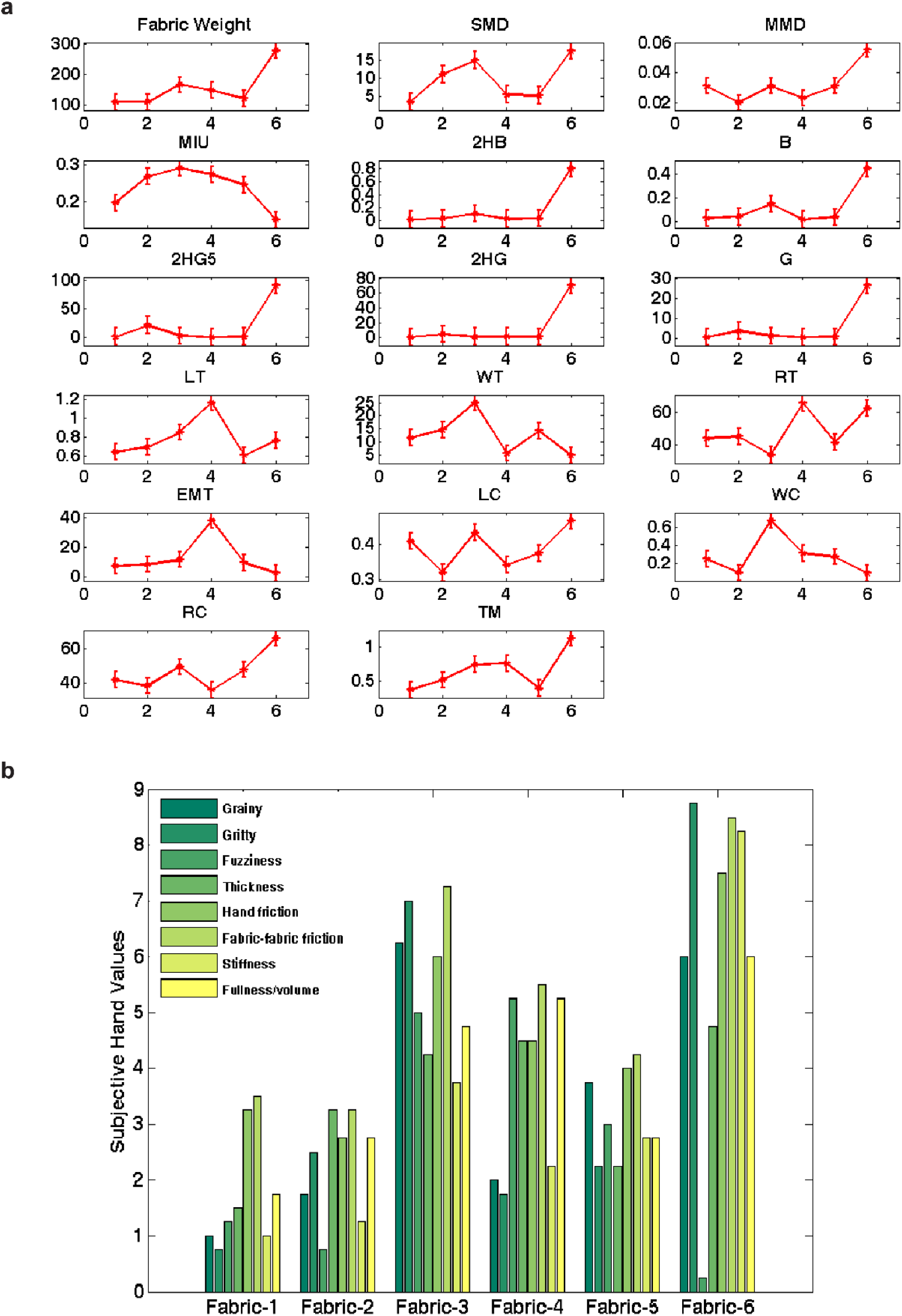
(a) Mean values of the low stress mechanical properties of the fabrics measured with KES and (b) Subjective hand evaluations of the fabrics. The bar graph shows the mean subjective evaluations of the six fabrics on a 0-9 Likert scale

### Subjective Evaluations

Subjective hand evaluations rated by the twenty participants were averaged, and the mean values are shown in Figure-2b. Surface roughness characteristics (grainy and gritty) were rated high for F3 and F6. F1 received the lowest rating of all in those two hand measures. Hand and fabric-fabric friction were also rated high on F3 and F6 and low for fabrics F1 and F2. Stiffness was rated highest in F6 and lowest in F1. Fabrics F3 and F4 rated higher for fuzziness. Regarding thickness and fullness/volume, F6 was rated the highest, and the lowest was F1.

### Biosignal Characteristics

The temporal profiles of the EEG and the GSR signals were computed to determine if they were distinguishable for the different fabric samples. In order to estimate the temporal changes in the EEG signal induced by fabric stimuli, the acquired signal was segmented into regions of contiguous sections in which an external stimulus is present, preceded by the absence of a stimulus. The fabrics showed a visibly varying EEG spectrum with activities in the range of 2 to 16 Hz, which can be seen in Figure-3a, where the spectrogram of one subject was plotted for the six fabrics. To study the tactile sensory responses, the average temporal profile of EEG signals at α-band (8-14 Hz) was extracted (see Figure-3b) for each fabric using all the subjects’ data. The fabrics stimulated distinguishable time-varying amplitude features in the EEG signal. Figure-3c shows the probability distribution of EEG amplitudes for all six fabrics, limited to the α-band. Kolmogorov–Smirnov (KS) test showed a significant difference (α = 0.05, p<.001) in the EEG amplitude distribution among the fabrics tested, except F3 vs. F5 (α = 0.05, p = .197). Fabric samples F3 and F4 induced higher EEG signal amplitudes than the rest of the fabrics, and F1 showed the lowest. The plot demonstrates that the neural responses differ for each fabric tested here.

**Figure 3.**
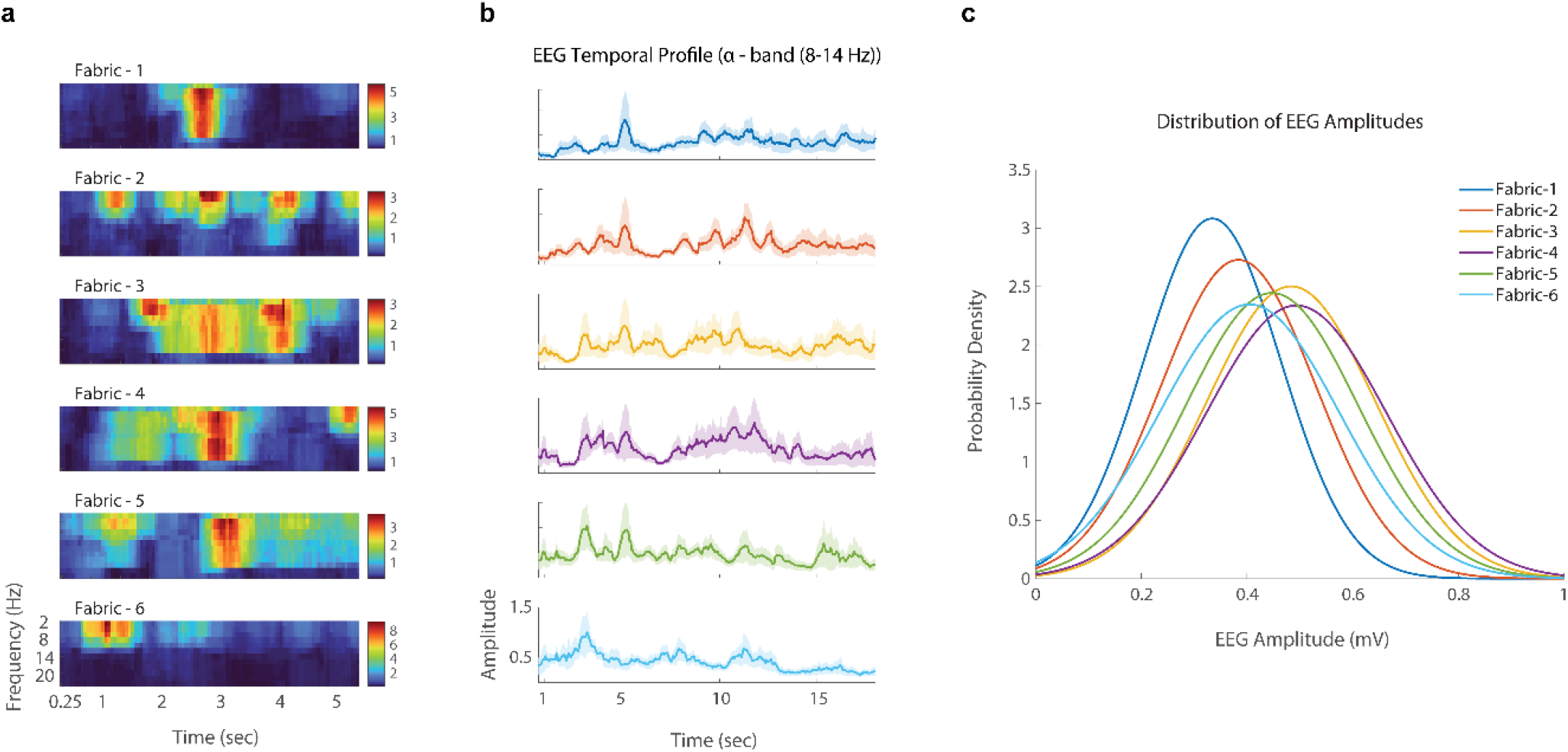
(a) Specimen EEG spectrogram of one subject for all six fabrics. The regions of red indicate high activation and blue corresponds to the lowest activation (b) Mean temporal profile of EEG signal at alpha frequency (8-14 Hz) range for each fabric (c) Probability distribution of EEG amplitudes (αEEG band) for all the six fabrics

The GSR signals were analyzed to estimate (i) the amplitude of skin conductance fluctuations (aGSR) and (ii) the number of skin conductance fluctuations (nGSR). nGSR refers to the number of reversals occurring per unit of time, and aGSR is the root-mean-square value of the skin conductivity signals measured over a unit of time. Figure-4 shows (a) a sample skin conductance signal of one subject for all six fabrics, (b) GSR amplitude distribution, and (c) the average number of skin conductance fluctuations. Overall, the fabric samples F1, F2, F3, and F5 induced higher electrodermal response measured as nGSR than samples F4 and F6. Similar to EEG signal distribution, a KS test was performed on the aGSR signals. F6 showed a significant difference from the rest of the fabrics, F1, F2, and F3 did not show significant differences among themselves but differed from the rest, and F4 and F5 differed significantly. The criterion for significance was the same as the EEG, i.e., α = 0.05, p<.001.

**Figure 4.**
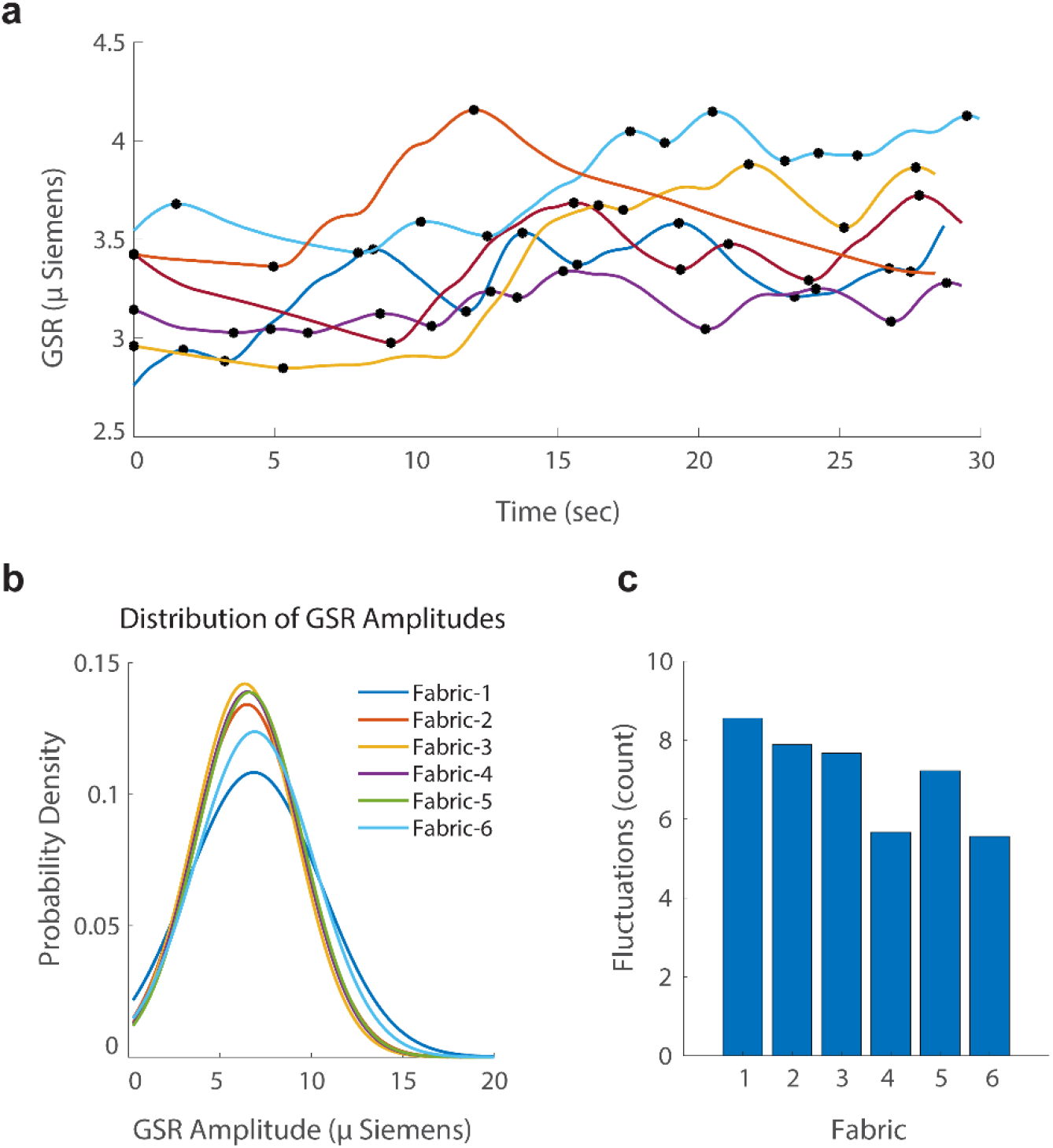
(a) Specimen GSR signal from one subject for all six fabrics; the marked dots represent the reversal points (b) Probability distribution of GSR amplitudes (aGSR) and (c) Average number of GSR fluctuations (nGSR) for all the six fabrics

### Correlation Analyses

To further explore the influence of the individual fabric properties on the biosignals, correlation analysis using Spearman’s rho was computed for four pairs – (1) spectral power of αEEG signals and human subjective hand evaluation, (2) spectral power of αEEG signals and LSM properties of the fabrics, (3) nGSR and human subjective hand evaluation and (4) nGSR and LSM properties of the fabrics. The LSM properties were classified into Tensile, Compression, Bending & Shear, and Friction & weight categories for convenience.

### αEEG

The correlation between the αEEG and the human subjective evaluations is shown in Figure-5a. Correlations with the LSM properties are shown in Figure-5b. All eight subjective ratings showed negative correlations with the αEEG signal power. Thickness and Fullness/Volume showed strong inverse correlations, with corresponding rho values of −0.62 and −0.57. Grainy and stiffness showed a moderate correlation with −0.47 and −0.42, respectively. Fuzziness showed the least correlation with αEEG with rho of −0.25

In the case of the LSM properties, the association strength varied from 0.11 to -.54. All properties, except LC, showed a negative correlation with the αEEG spectral power. The correlation of tensile properties (LT, EMT, RT, WT) was in the range of −0.10 to -.33. In the case of bending and shearing (B, 2HG5, 2HB, G, 2HG), the range was from -.22 to -.29 and in the case of friction and weight (SMD, WT, MIU, MMD), the range was from −0.06 to -.54. Finally, for the compression (TM, RC, WC, LC) correlation was noted for TM, RC, WC with a range of -.11 to.-.49 and a positive correlation for LC (0.10). The results showed that out of the 17 LSM properties, SMD and TM had a strong association with neural activity, and others had a weak association (See Figure 5b).

**Figure 5.**
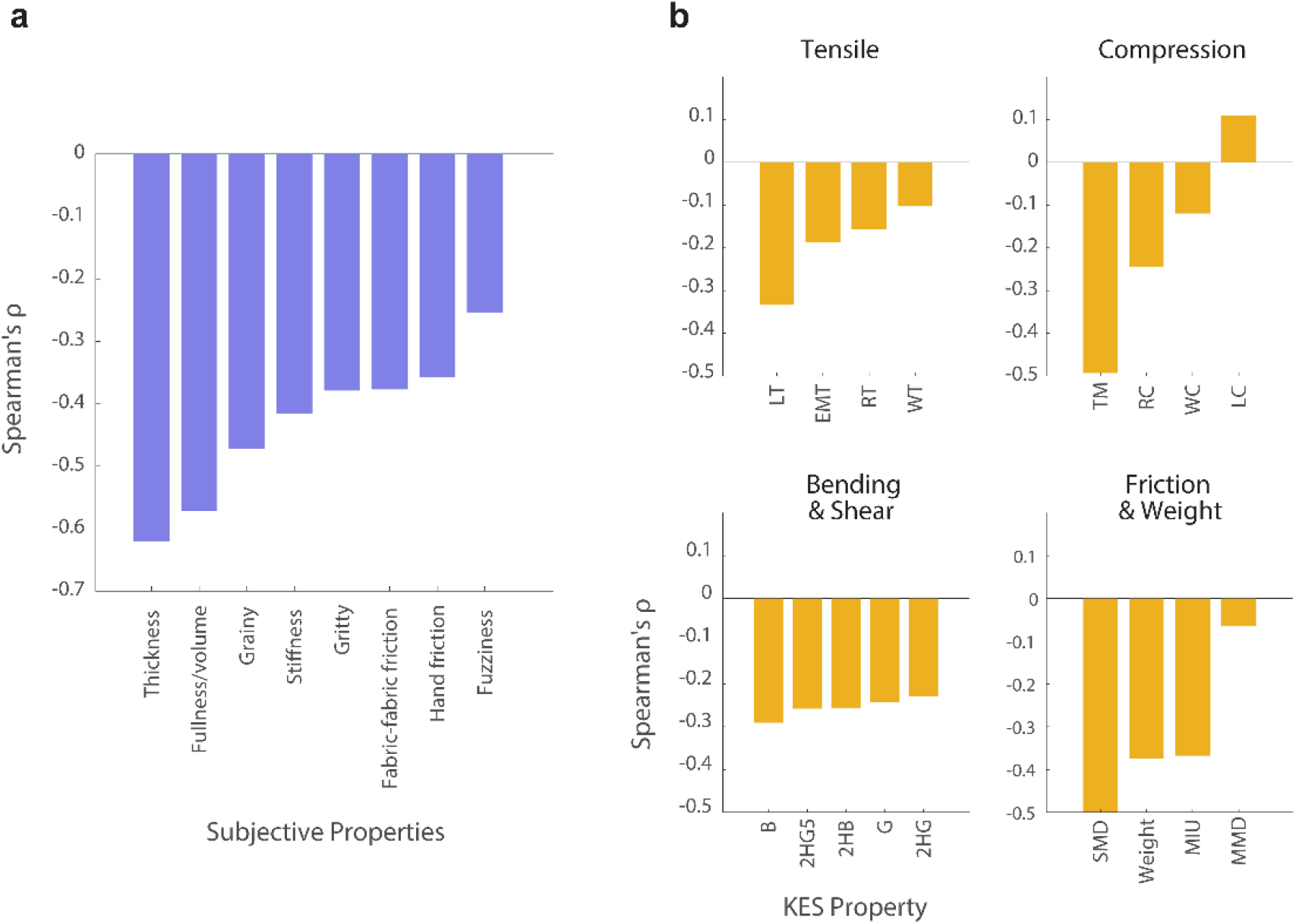
(a) Correlation estimates between αEEG spectral power and subjective hand evaluations and (b) Correlation between αEEG spectral power and LSM properties

### nGSR

The nGSR was correlated with the fabrics’ subjective evaluations and LSM properties. The nGSR showed a strong negative relationship with the subjective evaluations, with rho values ranging from −0.86 to −0.17 (see Figure 6a). Fullness/Volume showed the highest inverse correlation (−0.86), followed by thickness (−0.76), stiffness (−0.65), fabric-fabric friction (−0.63), and hand friction (−0.62). Like the αEEG, the nGSR showed a weak correlation with the fuzziness property.

**Figure 6.**
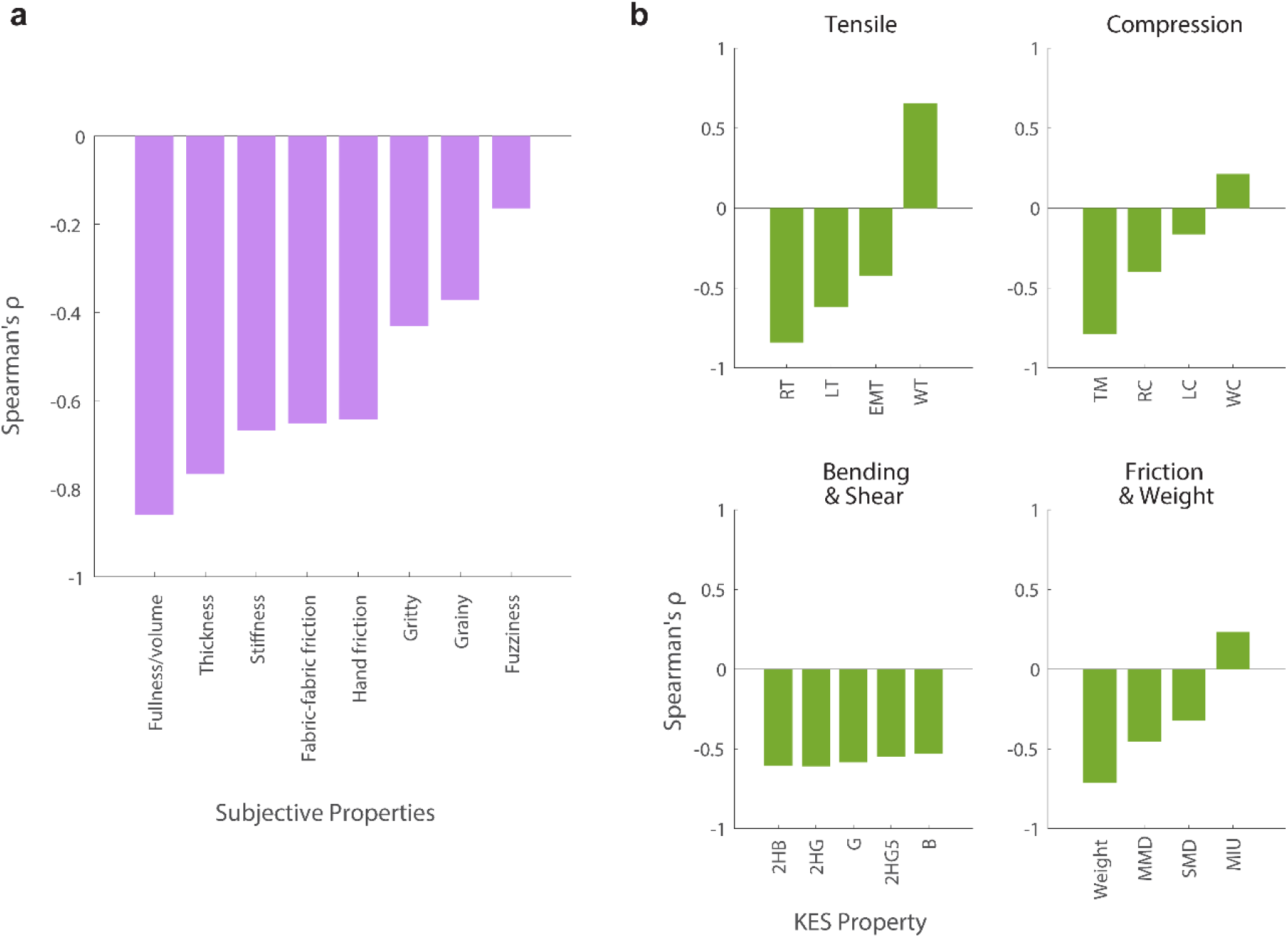
(a) Correlation between nGSR and subjective hand evaluation and (b) Correlation between nGSR and LSM properties

In the case of LSM properties versus nGSR, the association strength varied from 0.65 to -.84 (see Figure 6b). Three tensile properties (LT, EMT, RT) showed negative correlation with the nGSR with a range of −0.42 to −0.84 and the tensile energy (WT) showed a positive correlation with a strength of 0.65. In the case of bending and shearing (B, 2HG5, 2HB, G, 2HG), the range was from −0.53 to −0.60, and in the case of friction and weight (SMD, MIU, MMD), the range was from 0.25 to −0.45. Among the friction & weight category, the weight showed a strong positive correlation with nGSR (0.7). Finally, for the compression (TM, RC, WC, LC) negative correlation was found for TM, RC, LC with a range of -.16 to.-.78 and a positive correlation for WC (0.21).

## Discussion

Tactile sensation is a complex phenomenon that involves afferent pathways bringing sensory information into the brain, where the sensory cortex processes the information to generate perception. Most of the studies have focused on understanding this from the perspective of the brain in terms of areas that are involved in the process (Rajaei *et al*., 2018; Wang *et al*., 2016, 2019), how the sensory cortex perceives smoothness or roughness (Hoefer *et al*., 2016; Tang, Zhang, *et al*., 2022), how different fabric types can induce different brain activity (Rajaei *et al*., 2018; Yuan, Yu, Wang, *et al*., 2019) and how different textures elicit distinguishable activation patterns in the brain (Henderson *et al*., 2022; Moungou *et al*., 2015; Tang, Shu, *et al*., 2022). In this study, we explored tactile sensation from the perspective of fabric properties. We attempted to answer the question, what are fabrics’ primary features or properties that influence the human perception during active touch? And can we order them according to their influence?

During natural tactile interaction with fabrics, humans use their palm area and digits to actively feel the material without delineating the individual fabric properties. Such interaction is referred to as active touch (Delis *et al*., 2018). At the sensory cortex, during active touch, neural activity occurs in a broad spectral range (alpha, beta, theta, and delta) that are measurable using surface EEG electrodes. However, we focused on the alpha band in this study due to its relevance to attention to tactile features. Whenever more attention is needed to process tactile information, the alpha band attenuates, i.e., its power decreases. This behavior of the alpha band serves as the basis to determine which features of the fabrics generate more tactile information that the sensory cortex finds important and responsive to. Another feature that is responsive to tactile stimuli is the GSR. The relationship between tactile sensation and GSR at the palmer/planter regions has been studied since the 1930s in humans and non-human primates (Bach *et al*., 2010; Boucsein, 2012; Cardini and Haggard, 2013; Gatti *et al*., 2018; Turpin and Siddle, 1979). GSR-related studies postulate at least three important theories about the nature of the response, i.e. (i) the direct reflection of bio-electric muscle activity on skin conductivity, (ii) the connections with vasodilatation or vasoconstriction and (iii) its reflection on the electrical activity of the sweating glands (McCleary, 1950). During active touch stimulation, the skin conductance increases and fluctuates as one or more combinations of these phenomena occur. Hence, nGSR and aGSR can be used as indicators for evaluating sensory stimulations. To date, GSR responses have received less attention compared to EEG activities. Therefore, we attempted to use both amplitude and the number of reversals that occur during tactile stimulation, as they both are found to be responsive to touch. Though it fluctuated with active touch, the GSR amplitude was not distinguishable for individual fabrics presented. The number of reversals has been studied as a reliable metric indicating human physiological and psychological/emotional perception (Cardini and Haggard, 2013). In this study, the number of reversals in the GSR activity proved to be an effective metric for studying tactile stimulus from various fabric properties.

By individually correlating the eight subjective properties and the 17 LSM properties of the six fabrics with the αEEG spectral power, we were able to estimate how strongly each of the properties influenced the EEG activity at the sensory cortex. Likewise, the fabric properties were also correlated against the nGSR. Both αEEG and nGSR showed negative correlations with all the subjective properties, indicating that the inverse relation caused αEEG power to drop and the GSR to maintain its magnitude without fluctuating as the fabric properties increased in their magnitude. Thickness and Fullness/Volume were the dominant features influencing both biosignals. The friction and stiffness-related properties followed in their dominance, with Fuzziness being the least influencing property. Similar results were found in the cases of LSM properties. Thickness, weight, linearity of tensile force, and surface friction properties were the features that caused the αEEG to drop the most, as well as minimizing the nGSR. In both the biosignals, bending & shear properties showed a moderate relationship and were more or less equal for bending and shear forces and hysteresis values. Linearity of compression was the only property from the LSM that positively correlated with the αEEG signals. However, the correlation was weak. The three LSM properties that showed a positive correlation with the nGSR were the Tensile Energy, Compression Energy, and the Coefficient of surface friction; however, the latter two were weakly correlated.

The presence of correlation may not establish a causal relationship between these biosignals and the fabric properties. However, we can cautiously argue that the fabric properties have a causal influence on these signals. Furthermore, any non-linear relationship between the properties and the biosignals was not studied here. Due to the various layers of sensory transfer functions present in the human body, that alter the stimuli before they reach the sensory cortex, a linear relationship may not be considered thorough. By building simulation models to replicate cortical sensory signals, these non-linear transfer functions can be systematically studied in the future.

## Conclusion

Fabric properties are perceived by humans during touch. These perceptions often impart their signatures in neural and other bodily signals. By measuring the EEG and the GSR during active touch, the correlation between the individual fabric properties and the elicited responses were studied. We have shown that not all the fabric properties are equally perceived by the human sensory system. Specific fabric properties dominate the perceived sensations.

